# Benchmarking Large Language Models for Predicting Therapeutic Antisense Oligonucleotide Efficacy

**DOI:** 10.64898/2026.02.17.706455

**Authors:** Abhinit Sundar, Zhi Wei, Stephen Griesmer

## Abstract

Antisense oligonucleotides (ASOs) are a promising class of therapeutic drugs that can target and modulate genes associated with various diseases. This study benchmarks Large Language Models (LLMs) for predicting ASO therapeutic efficacy through a two-stage approach: (1) molecular embedding-based fine-tuning using SMILES representations, and (2) prompt engineering with zero-shot and few-shot learning using DNA sequences with target gene information. We evaluated general-purpose models (GPT-3.5-Turbo, LLaMA2-7B, Galactica-6.7B) and chemistry-specific models (ChemBERTa, Molformer, BERT) across three datasets: PFRED (522 sequences), openASO (1708 sequences), and ASOptimizer (1267 sequences). DNA sequence inputs with target gene information outperformed SMILES representations. GPT-3.5-Turbo achieved R^2^ values of 0.6381 (PFRED) and 0.6340 (ASOptimizer) for few-shot prompting with k=3 examples.

Code and datasets available at: https://github.com/asundar0128/IndependentStudy

## I. INTRODUCTION

Antisense oligonucleotides (ASOs) have demonstrated unique therapeutic potential through sequence-specific binding to target RNA sequences [7]. These molecules can modulate gene expression by targeting the transcription and translation processes, making them valuable tools for treating diseases at the genetic level.

Traditional ASO design approaches relied heavily on researcher expertise and physical observations, which became inadequate as the chemical space expanded exponentially with increasing RNA sequence diversity [8]. Each ASO position can be occupied by one of four nucleotide bases, leading to 4^*n*^ potential combinations for ASOs of length *n*. This vast combinatorial space necessitated computational approaches to efficiently screen ASO candidates. Early computational methods utilized linear models and thermodynamic calculations, but recent advances in machine learning offer new opportunities for ASO optimization.

## II. RELATED WORK

ASOptimizer integrates engineering of an ASO sequence and chemical engineering of the ASO to predict efficacy. Sequence engineering considers sequence complementarity as well as thermodynamic stability as measured by changes in Gibbs Free Energy changes with target binding vs. non-target binding [2]. The framework employed both sequence engineering (base composition optimization, motif selection) and chemical engineering (LNA modifications, phosphorothioate linkages) to enhance ASO efficacy.

PFRED (Pfizer Functional RNAi Enumeration and Design Tool) provides a comprehensive platform for ASO and small interfering ribonucleic acid (siRNA) design with GUI-based interaction and cloud deployment capabilities [1]. The tool incorporates bioinformatic descriptors like sequence position and composition, auto- and cross-covariance, and thermodynamic stability and flexibility into a support vector machine (SVM) model.

OpenASO is a research initiative to develop design principles for RNA-based therapeutics. A project under this initiative focused on identifying regulatory targets of an RNA molecule. A team, working under Lela Lackey, developed a support vector machine regression algorithm and ASO dataset for prediction of ASO efficacy [15]. OpenASO datasets contain ASO sequences with efficacy labels across multiple targets, focusing on biological dependencies between gene sequences and targets with various chemical modifications including phosphorothioate backbones and 2’-O-methyl groups.

Large Language Models (LLMs) have shown remarkable success in various domains, including chemistry and biology [3,4,5]. Recent advances in LLMs for molecular applications have demonstrated their potential for chemical property prediction [3,4]. Domain-specific models like ChemBERTa have been developed for molecular representation learning [12], while general-purpose models like GPT-3.5 [9] have shown versatility across diverse tasks through instruction tuning. This study investigates their potential for predicting ASO therapeutic efficacy through two distinct methodological stages, comparing their performance against traditional approaches and baseline models.

## III. METHODOLOGY

### A. Two-Stage Experimental Design

Our experimental approach consisted of two distinct stages to comprehensively evaluate LLM performance for ASO efficacy prediction:

#### Stage 1

Molecular embedding-based approach using SMILES representations converted from DNA sequences, followed by fine-tuning of chemistry-specific LLMs (ChemBERTa [12], Molformer [13], BERT [14]) with ridge regression for efficacy prediction.

#### Stage 2

Prompt engineering approach using DNA sequences with target gene information, evaluating general-purpose LLMs (GPT-3.5-Turbo [9], LLaMA2-7B [10], Galactica-6.7B [11]) through zero-shot and few-shot learning paradigms. Zero-shot learning involves model prediction without any example ASO-efficacy pairs in the prompt, while few-shot learning provides k=3 example ASO sequences with their known efficacy values to guide the model’s predictions.

### B. Dataset Characteristics

Three datasets were utilized with established baselines derived from their respective computational frameworks:

- **PFRED (522 sequences):** Baseline R^2^ = 0.28 from thermodynamic stability calculations using OligoWalk [1]
- **openASO (1708 sequences):** Baseline R^2^ = 0.3028 from transcript-level binding predictions [15]
- **ASOptimizer (1267 sequences):** Baseline R^2^ = 0.4020 from integrated Gibbs Free Energy and MIRANDA target prediction models [2]

### C. Model Selection and Evaluation

Chemistry-specific models included ChemBERTa (transformer-based molecular representation) [12], Molformer (molecular transformer) [13], and BERT (adapted for chemical sequences) [14]. General-purpose models comprised GPT-3.5-Turbo (instruction-tuned for diverse tasks) [9], LLaMA2-7B (conversational AI) [10], and Galactica-6.7B (scientific reasoning) [11].

Few-shot experiments used exactly k=3 examples randomly selected from training data for all tests to ensure consistency. Evaluation metrics included Root Mean Square Error (RMSE) and coefficient of determination (R^2^). Models beyond GPT-3.5 showed inferior performance and are not reported in detail.

## IV. EXPERIMENTAL RESULTS

### A. Stage 1: Molecular Embedding Approach

Stage 1 results demonstrate that Molformer achieved the highest R^2^ values for PFRED (0.3072) and ASOptimizer (0.3774) datasets, while BERT performed best for openASO (R^2^ = 0.2231). Analysis of both RMSE and R^2^ metrics reveals that most models underperformed compared to baseline metrics, indicating limitations of SMILES-based molecular embeddings for capturing ASO-specific biological interactions. Missing RMSE values in the ASOptimizer column are due to computational resource constraints during model training.

**TABLE 1.** STAGE 1 RESULTS: MOLECULAR EMBEDDING APPROACH.

| Model | PFRED |  | openASO |  | ASOptimizer |  |
| --- | --- | --- | --- | --- | --- | --- |
| | RMSE | $R^2$ | RMSE | $R^2$ | RMSE | $R^2$ |
| Baseline | 0.2422 | 0.28 | 0.2283 | 0.3028 | 0.2091 | 0.4020 |
| ChemBERTa | 0.3252 | 0.1181 | 0.2938 | 0.0623 | — | 0.2414 |
| BERT | 0.3162 | 0.0570 | 0.3153 | 0.2231 | — | 0.2761 |
| Molformer | 0.5021 | 0.3072 | 0.3038 | 0.1384 | — | 0.3774 |
| Mol2Vec | 0.3302 | 0.0328 | 0.2769 | 0.0135 | — | 0.0126 |

### B. Stage 2: Prompt Engineering Approach

Stage 2 results reveal significant performance differences between zero-shot and few-shot approaches. Zero-shot prompting involves the model making predictions based solely on its pre-trained knowledge without example ASO-efficacy pairs, while few-shot prompting with k=3 provides three example ASO sequences with known efficacy values to guide predictions. GPT-3.5-Turbo demonstrated superior performance on PFRED and ASOptimizer datasets when comparing zero-shot (no examples) versus few-shot (k=3 examples) approaches, with few-shot learning showing substantial improvements in R^2^ values. However, the openASO dataset presented unique challenges, with all models showing negative R^2^ values, indicating performance worse than a naive mean predictor.

**TABLE 2.** STAGE 2 RESULTS: PROMPT ENGINEERING APPROACH.

| Model | Zero-Shot | | Few-Shot ($k=3$) | |
| --- | --- | --- | --- | --- |
| | RMSE | $R^2$ | RMSE | $R^2$ |
| PFRED Dataset |  |  |  |  |
| Galactica-6.7B | 0.4745 | -1.4528 | 0.4302 | -1.3388 |
| LLaMA2-7B | 0.4286 | -1.3389 | 0.4212 | -1.2588 |
| GPT-3.5-Turbo | 0.3275 | 0.3637 | 0.3591 | **0.6381** |
| openASO Dataset |  |  |  |  |
| Galactica-6.7B | 0.5925 | -3.6965 | 0.3515 | -0.6530 |
| LLaMA2-7B | 0.4437 | -1.6328 | 0.3570 | -1.0832 |
| GPT-3.5-Turbo | 0.2822 | -0.0651 | 0.3248 | -0.4094 |
| ASOptimizer Dataset |  |  |  |  |
| Galactica-6.7B | 0.4478 | -1.7443 | 0.3813 | -0.9888 |
| LLaMA2-7B | 0.4382 | -1.6310 | 0.4080 | -1.2078 |
| GPT-3.5-Turbo | 0.3185 | 0.3471 | 0.3506 | **0.6340** |

All few-shot experiments used exactly $k=3$ examples for consistency across datasets and models.

## V. DISCUSSION

### A. Performance Analysis

The performance of DNA sequence-based inputs with target gene information compared to SMILES representations highlights the importance of biological context in ASO efficacy prediction [8]. DNA sequences with target gene information provide sequence-function relationships that SMILES cannot capture, as demonstrated by the superior performance of Stage 2 approaches over Stage 1 molecular embeddings.

GPT-3.5-Turbo’s performance stems from its instruction-tuning and reasoning capabilities [9], allowing interpretation of biological contexts without domain-specific fine-tuning. The model generated more diverse predictions compared to other models which showed clustering behaviors around specific values.

### B. Dataset-Specific Challenges

The openASO dataset’s poor performance across all models (negative R^2^ values) suggests it may contain more complex sequence-target relationships or experimental noise that current LLM approaches cannot model effectively. This contrasts with the success on PFRED and ASOptimizer datasets, indicating that certain ASO characteristics or experimental protocols may be more amenable to LLM-based prediction.

### C. Future Improvements

Dataset expansion to include more gene targets, chemical modifications, and experimental conditions would provide richer training signals. Enhanced prompting strategies incorporating biochemical principles and chain-of-thought reasoning may improve prediction accuracy. Hybrid approaches combining molecular embeddings with prompt engineering should be explored.

## VI. CONCLUSION

This study demonstrates that LLMs show promise for therapeutic ASO efficacy prediction, with DNA sequence-based approaches incorporating target gene information outperforming SMILES-based molecular embeddings. GPT-3.5-Turbo achieved notable R^2^ improvements over baselines on PFRED and ASOptimizer datasets. The universal failure on openASO highlights important limitations requiring further investigation.

Future research should focus on hybrid methodologies, specialized fine-tuning approaches, and understanding dataset-specific factors that influence LLM performance in ASO design applications.

## Addendum: Reproducibility, Experimental Clarification, and Limitations

### Experimental Configuration and Reproducibility

All experiments were conducted using Python 3.10 and scikit-learn. Few-shot prompt examples were randomly selected from training data using a fixed random seed to ensure reproducibility. RMSE and R^2^ metrics were computed using standard scikit-learn implementations on held-out test sets. All code and evaluation scripts are publicly available at the provided GitHub repository.

### Experimental Scope Clarification

Few-shot prompting experiments used k=3 examples to balance prompt efficiency with predictive performance while maintaining consistent experimental conditions across models and datasets.

### Limitations

Few-shot performance depends on prompt example selection and prompt design. General-purpose LLMs are not specifically trained on ASO datasets, which may limit biological specificity. These results represent computational benchmarking and do not constitute experimental or clinical validation.

## REFERENCES

[1] S. Sciabola et al., “PFRED: A computational platform for siRNA and antisense oligonucleotides design,” PLoS One, vol. 16, no. 1, article e0238753, Jan. 2021.

[2] G. Hwang et al., “ASOptimizer: Optimizing Antisense Oligonucleotides Through Deep Learning for IDO1 Gene Regulation,” Molecular Therapy: Nucleic Acids, vol. 35, June 2024.

[3] T. Guo et al., “What Can Large Language Models Do in Chemistry? A Comprehensive Benchmark on Eight Tasks,” Proc. 37th Conf. Neural Information Processing Systems, 2023.

[4] S. Sadeghi et al., “Can large language models understand molecules?” BMC Bioinformatics, vol. 25, no. 1, article 225, June 2024.

[5] X. Tang et al., “MolLM: A Unified Language Model for Integrating Biomedical Text with 2D and 3D Molecular Representations,” Bioinformatics, vol. 40, suppl. 1, pp. i357–i368, 2024.

[6] J. M. Zambrano Chaves et al., “Tx-LLM: A Large Language Model for Therapeutics,” arXiv, June 2024.

[7] S. T. Crooke et al., “Antisense Technology: A Review,” Journal of Biological Chemistry, vol. 296, article 100416, 2021.

[8] P. H. Hagedorn et al., “Managing the sequence-specificity of antisense oligonucleotides in drug discovery,” Nucleic Acids Research, vol. 45, no. 5, pp. 2262–2282, Mar. 2017.

[9] OpenAI, “GPT-4 Technical Report,” arXiv preprint arXiv:2303.08774, Mar. 2023.

[10] H. Touvron et al., “Llama 2: Open Foundation and Fine-Tuned Chat Models,” arXiv preprint arXiv:2307.09288, July 2023.

[11] R. Taylor et al., “Galactica: A Large Language Model for Science,” arXiv preprint arXiv:2211.09085, Nov. 2022.

[12] S. Chithrananda et al., “ChemBERTa: Large-Scale Self-Supervised Pretraining for Molecular Property Prediction,” arXiv preprint arXiv:2010.09885, Oct. 2020.

[13] J. Ross et al., “Large-scale chemical language representations capture molecular structure and properties,” Nature Machine Intelligence, vol. 4, pp. 1256–1264, Dec. 2022.

[14] J. Devlin et al., “BERT: Pre-training of Deep Bidirectional Transformers for Language Understanding,” Proc. 2019 Conf. North American Chapter of the Association for Computational Linguistics, pp. 4171–4186, 2019.

[15] L. Lackey et al., “OpenASO: A machine learning approach for antisense oligonucleotide design,” GitHub repository. Available: https://github.com/Genentech/OpenASO

